# A scalable open-source framework for machine learning based image collection, annotation and classification: a case study for automatic fish species identification

**DOI:** 10.1101/2022.06.29.498112

**Authors:** Catarina NS Silva, Justas Dainys, Sean Simmons, Vincentas Vienožinskis, Asta Audzijonyte

## Abstract

Citizen science platforms, social media and multiple smart phone applications enable collection of large amounts of georeferenced images. This provides a huge opportunity in biodiversity and ecological research, but also creates challenges for efficient data handling and processing. Recreational and small-scale fisheries is one of the fields that could be revolutionised by efficient, widely accessible and machine learning based processing of georeferenced images. The majority of non-commercial inland and coastal fisheries are considered data poor and are rarely assessed, yet they provide multiple societal benefits and can have large ecological impacts. Given that large quantities of fish observations and images are being collected by fishers every day, artificial intelligence (AI) and computer vision applications offer a great opportunity to improve data collection, automate analyses and inform management. Yet, to date, many AI image analysis applications in fisheries are focused on the commercial sector and are not publicly available for community use. In this study we present an open-source modular framework for large scale image storage, handling, annotation and automatic classification, using cost- and labour-efficient methodologies. The tool is based on TensorFlow Lite Model Maker library and includes data augmentation and transfer learning techniques, applied to different convolutional neural network models. We demonstrate the implementation of this framework in an example case study for automatic fish species identification from images taken through a recreational fishing smartphone application. The framework presented here is highly customisable for further advancement and community based image collection and annotation.

## 1. Introduction

More than 80% of global catches occur in fisheries that lack essential data, resources, and infrastructure for stock assessments to be performed (Costello *et al*. 2020). This is especially true for recreational fisheries, which, in the developed world at least, continue to grow in popularity and have important well-being and economic benefits, but remain hard to monitor, control and assess (Meirelles *et al*. 2020).

Given the large number of people engaged in recreational fisheries and generally high level of technology used, there is a potential for large scale data collection that could greatly improve our knowledge about recreational catches and the populations’ status. There is generally a strong motivation among recreational fishers to conserve fish stocks and experience with other groups have shown that engagement in citizen science programs does not only help to generate large datasets, but also promote awareness and sense of stewardship (Dickinson *et al*. 2010). For recreational fisheries management, citizen science would be especially powerful because it could enable collaborative research and management which has been shown to have clear benefits across the world (Venturelli *et al*. 2017; Harris *et al*. 2021). Therefore, it is urgently due that recreational fisheries management benefits from the increasing popularity of mobile fishing applications to achieve a step-change in data collection and angler engagement.

Artificial intelligence (AI) has already revolutionised weather forecasting, wildfires disaster response, health care and transportation. AI and computer vision applications also offer a great untapped opportunity to transform recreational fisheries management because they allow rapid processing of large citizen science datasets, including automation of species identification and potentially also the fish size measurement. Even though research applying AI in fisheries has been increasing, with about 40 scientific publications per year (Ebrahimi *et al*. 2021), this is still very limited compared to other fields and mainly applied to commercial fisheries (e.g. Lekunberri *et al*. 2022; Ovalle *et al*. 2022). Moreover, the methods, tools and scripts developed in these studies are often not publicly available, limiting wider uptake, application and community-driven improvement.

To help address the issue of limited AI application in recreational and small-scale fisheries research and management we present a modular open source framework for management and visual recognition of large image collections. The framework includes steps for 1) data management (storage and pre-processing), 2) image processing (automatic detection of fishes from images with pre-trained models, manual annotation of species supported by metadata and images augmentation) and 3) machine learning model development (train and test algorithms for species classification and detection). Alongside the framework, we also summarise currently available open-source tools and provide scripts that can be customised by researchers and applied to different types of imagery data. We demonstrate the implementation of the framework and its potential use for recreational fisheries research, through a pilot study that aims to automate detection of fish species from images uploaded to a smartphone fishing application. Finally, although this framework is developed for fisheries, it could also be applied to other areas that require image annotation, processing and classification.

## 2. Framework

The framework developed in this study is summarised in figure 1 and is divided into three main modules: data management, image processing and machine learning. This framework has a variety of applications and the different components and scripts can be customized to different image classification oriented projects and used for example for species/individual identification, size estimation and other phenotypic/morphological pattern identification from images.

**Figure 1:**
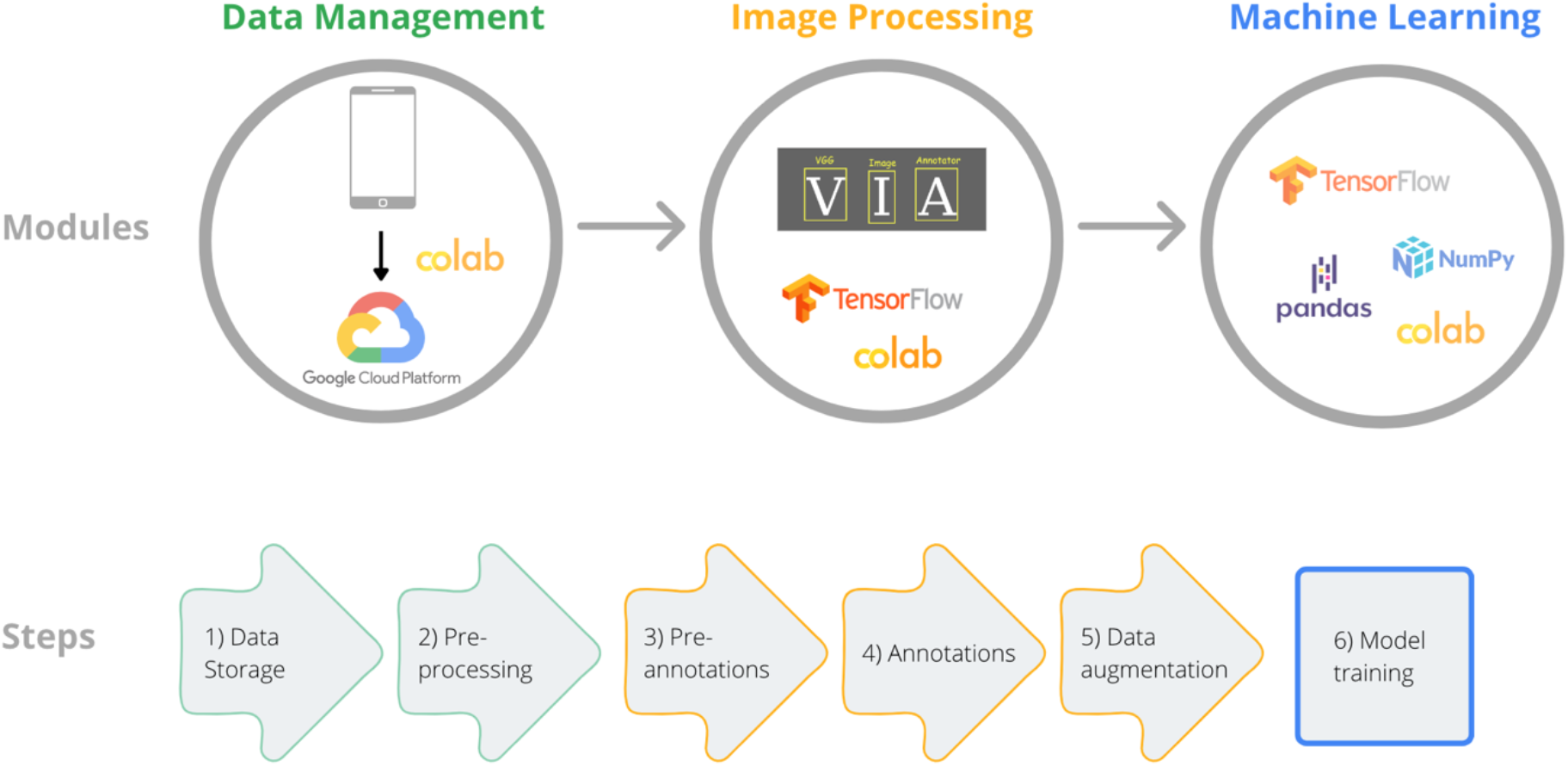
Overview of the framework, with the main tools used, consisting of three modules: and six steps. Platforms and specific tools (e.g. Google colab and Google Cloud Platform) indicated here were used in the example application, but could be replaced with other tools, as explained below. All steps described are supported by a freely available code library, deposited in https://github.com/FishSizeProject

Computer vision is a branch of computer science which aims to extract information from images (for example from photos and movies; Prince 2012) and develop visual recognition systems. Some of the most commonly used methods are image classification and object detection. Image classification is a technique used to classify or predict the class of a single object in an image (i.e., single-label classification; Mohri *et al*. 2012). Object detection is used to detect the location of one or more objects in a given image and then categorise each object (i.e. predict the class of each object, also called object classification). Object detection can be achieved by annotating images with a rectangle or bounding box around the object (see for example dos Santos & Gonçalves 2019).

### Module 1: Data Management

The framework presented here assumes that users already have acquired the images. These images could have been provided by citizen science programs, social media scans or targeted image collection.

#### Step 1: Storage

The images and associated metadata are stored on remote servers, i.e. cloud storage. There are a number of benefits to cloud storage of large datasets, including costs, facilitated collaboration, easy access from multiple devices, access to virtual servers used for analyses (see below), efficient back-up, centralization and data protection. Given that many projects for image analyses may only require storage for a short period, the costs associated with cloud-based storage are likely to be lower than investing into local devices and hard drives. From our experience, storage services from the three main cloud storage providers are very similar and differ mostly in terminology (Table 1). Pricing structure and billing depend on the type of resource needed and is briefly discussed in the “Lessons learned” section. To access these services users only need a platform specific account and a payment method. Services typically provide a free trial for a limited amount of data and time which varies between providers (for example both Google Cloud Platform and Microsoft Azure Platform provide $200 credit to use in 30 days in any service and one of Amazon Web Services include 250 hours per month to use the ml.t3.medium computer instance).

**Table 1:** A comparison of the main storage services providers Amazon Web Services (AWS), Google Cloud Platform (GCP) and Microsoft Azure Platform (Azure) with specific terminology used by each provider.

| Service | AWS | GCP | Azure |
| --- | --- | --- | --- |
| Versioning | ✓ | ✓ | ✓ |
| Encryption | ✓ | ✓ | ✓ |
| Fine-grained security (multiple criteria can be used for authorization to access data/resources) | ✓ | ✓ | ✓ |
| Lower fees for less frequently accessed data | Infrequently accessed class | Cool Blob (binary large object) | Nearline (frequently accessed), Coldline (infrequently accessed) |
| Archiving | Glacier | Archive | Archive |
| Free tier service (use the product for free up to specified limits) | ✓ | ✓ | ✓ |
| On-demand charges for resources used | ✓ | ✓ | ✓ |

#### Step 2: Pre-processing – sensitive data protection

For all further steps in this study we used Google Colab (Bisong 2019), an online tool with built-in (i.e. requires no setup) interactive Python programming environments such as Jupyter notebooks (Kluyver *et al*. 2016) and with free access to computing resources such as GPUs (graphics processing units, which are essential for computer vision tasks and image processing). However, this could be replaced with other tools such as JupyterLab or Kaggle.

In many cases, images collected by citizens or extracted from internet may contain sensitive data, such as people’s faces. Depending on the nature of subsequent work (e.g. crowdsourced annotation of images), it may be preferrable to remove such data. In this framework we introduce a step that uses face detection algorithms to remove sensitive data before further analyses. This is done using the publicly available script *face_detection_overlay*.*ipynb* as a part of the general framework code. This script uses the function *detect_face()* of the Python library CVlib and the pre-trained model *caffemodel* to detect human faces (Ponnusamy 2018).

### Module 2: Image annotation

Different computer vision techniques require different types of annotations of images to be used for training and testing the models. Image annotations are often done manually and this step is frequently identified as one of the main bottlenecks for using machine learning approaches. This is particularly true in areas where image annotation for model training requires expert knowledge such as accurate identification of fish species. In this framework we focus on image classification and object detection using bounding boxes and do not include other computer vision tools such as image segmentation. This is because they are the least time and therefore resource consuming methods, which was an important criterion given the focus of our study on developing tools for research groups with limited resources.

#### Step 3: Pre-annotations – accelerate manual annotation of images

Although expert based manual annotation of images (into correct species or other groups that will be used in the model) cannot be avoided, there are several pre-annotation steps that can reduce the amount of manual work required. Specifically, our framework includes importing images from cloud storage, running an object detector using the module *inception_resnet_v2* a Keras image classification model pre-trained on Open Images Dataset V4 (Kuznetsova *et al*. 2020), converting the bounding boxes metadata to absolute coordinates and saving the metadata in VGG format (in a .csv file). The pre-trained object detector was used to place bounding boxes around all fish shapes in the image, but the model can be used to detect 600 shapes, including elephant, lynx, bird, insect, shellfish, tree, plant and others. The bounding box step is needed only for algorithms focused on the object detection method, not for image classification. If users are only interested in classification, the bounding box step can be skipped. However, automatic detection of specific shapes might still be useful if some images in the collection don’t have the object that needs to be classified. For example, the photos collected through angler apps may include pictures of location, gear or just accidental images. Running an object detection step will reduce the amount of work required to sort the images manually.

The scripts for step 3 are available in the notebook *object_detection_pre-annotation*.*ipynb*. The last section of the script includes formatting and saving bounding box coordinates in the input format required by the software used for manual annotations (VGG software, see below). This section of the script can be easily customised for other formats of annotation tools.

#### Step 4: Annotations – manual annotation of images

There is a variety of open-source software for manual image annotation (Table 2) with a range of formats for importing and exporting object annotations; each software package typically has its own Json-based or csv-based format. Some tools are only available online and therefore require uploading images to annotation servers. This may limit their application for sensitive data or in situations where experts engaged in image annotation have limited internet connectivity.

**Table 2:** List of open-source software for image annotation with details about export formats.

| Software | Export formats | Specifications | Web page |
| --- | --- | --- | --- |
| RectLabel | YOLO, Create ML, COCO json, csv | Runs locally; only available for MacOS operating system | https://rectlabel.com/ |
| Scalable | json | Runs locally; installation in terminal | https://scalabel.ai |
| VGG Image Annotator (via-2.0.11) | csv, VGG json, COCO | Runs locally; no installation or setup required; fast | https://gitlab.com/vgg/via |
| LabelImg | VOC xml, YOLO, createML | Runs locally; installation in terminal | https://github.com/tzutalin/labelImg |
| MakeSense | Yolo, VOC xml, VGG json, csv | Runs online but not functional for many images (if web browser resets, all annotations are lost) | https://www.makesense.ai/ |
| SuperAnnotate | json | Runs locally; easy installation (all operating systems) | https://www.superannotate.com |
| LabelBox | json | Runs online | https://labelbox.com/ |
| Supervisely | json | Runs online | https://supervise.ly/ |

In this study we used the VGG software (Dutta & Zisserman 2019), as it could be run locally and had easy setup and installation. The generated .csv file from the Step 3 (*object_detection_pre-annotation*.*ipynb*) was opened in the VGG software, where automatic pre-annotation of image shapes was inspected manually and corrected if needed (bounding boxes adjusted to better fit the object), and class names added. In our case class names included identification of fish species and this step required expert knowledge. If class names are provided automatically in the pre-annotation step (e.g. the model only aims to classify fishes or other shapes), they can also be manually corrected.

Because TensorFlow Lite Model Maker requires annotations input file in a specific format, we have also developed a script (*convert_annotations_VGG_to_TF*.*ipynb*) to format the .csv file from VGG software to the TensorFlow format. This includes converting bounding boxes coordinates from absolute values generated by the VGG software to relative values needed for TensorFlow and splitting the dataset into train, test and validation sets.

#### Step 5: Data augmentation

Data augmentation involves creating multiple copies of the same images, but with transformations such as flipping, rotating, scaling and cropping. Image augmentations have been shown to combat overfitting in deep convolutional neural networks (Shorten & Khoshgoftaar 2019), improve performance (Mikołajczyk & Grochowski 2018; Shorten & Khoshgoftaar 2019), model convergence (Liu *et al*. 2020), generalization and robustness on out-of-distribution samples (Bengio *et al*. 2011; Hendrycks *et al*. 2020), and, in general, to have more advantages compared to other methods (Hernández-García & König 2018). Depending on the method of computer vision used, data augmentation steps will differ. For example, for image classification data augmentation only involves transformations of the images. However, for the object detection method, when data augmentation is employed after annotations, as in this framework, augmentation also needs to be applied to the coordinates of bounding boxes (i.e. annotations need to be converted to be in agreement with image transformations).

In this framework we use the open source Albumentations library (Buslaev *et al*. 2020) for data augmentation. The script *data_augmentation_classification*.*ipynb* defines an augmentation pipeline for image classification approach and applies vertical and horizontal flips for all images in a directory. The script *data_augmentation_object_detection*.*ipynb* is used to transform the annotations (bounding boxes) for the object detection method by applying vertical and horizontal flip to the coordinates of the bounding boxes from the annotations file.

### Module 3: Machine learning

#### Step 6: Model training and testing

This framework uses the Tensorflow Lite Model Maker library (Abadi et al. 2016a; b) and transfer learning which reduces the amount of training data required and model training time. Tensorflow Lite supports several model architectures, including EfficientNet-Lite, MobileNetV2 and ResNet50 (He *et al*. 2016; Sandler *et al*. 2018; Tan & Le 2019) which are pre-trained models for image classification, and EfficientDet-Lite[0-4], a family of mobile and IoT-friendly models for object detection, derived from the EfficientDet architecture (Tan *et al*. 2020). The library is flexible and new pre-trained models can be added by customising the library code.

For this framework, we developed the script *image_classification*.*ipynb* to train and test (evaluate) an image classification model using the pre-trained models mentioned above. This script also generates a confusion matrix for visualizing model performance and functions to load a trained model and run classification inference on new images. The script *object_detection*.*ipynb* includes functions to train and test an object detection model.

## 3. Pilot case-study

To illustrate the feasibility of the framework developed here, we present a pilot case-study of detecting the species Common bream (*Abramis brama)*, European carp (*Cyprinus carpio*), Northern pike (*Esox lucius)*, Largemouth bass (*Micropterus salmoides)*, European perch (*Perca fluviatilis)* and Pikeperch (*Sander lucioperca)* (Figure 2). Images were obtained through a collaborative agreement with a company Fish Deeper™, which provides fishfinder devices which are popular among anglers and runs a smart phone application enabling anglers to log their catch. The anonymous data obtained included images and associated metadata, such as fish species identification by the user, GPS coordinates and other information. After the automated pre-annotation to select only images with fish, the manual image annotation was done by one person (Justas Dainys) with the required expertise in fish species identification (see discussion at the end for more details about the time used in this step). Next, we applied image augmentation (vertical and horizontal flips) to increase the number of images for model training, which provided 3 additional images for each original photo and resulted in a total of 4809 images.

**Figure 2:**
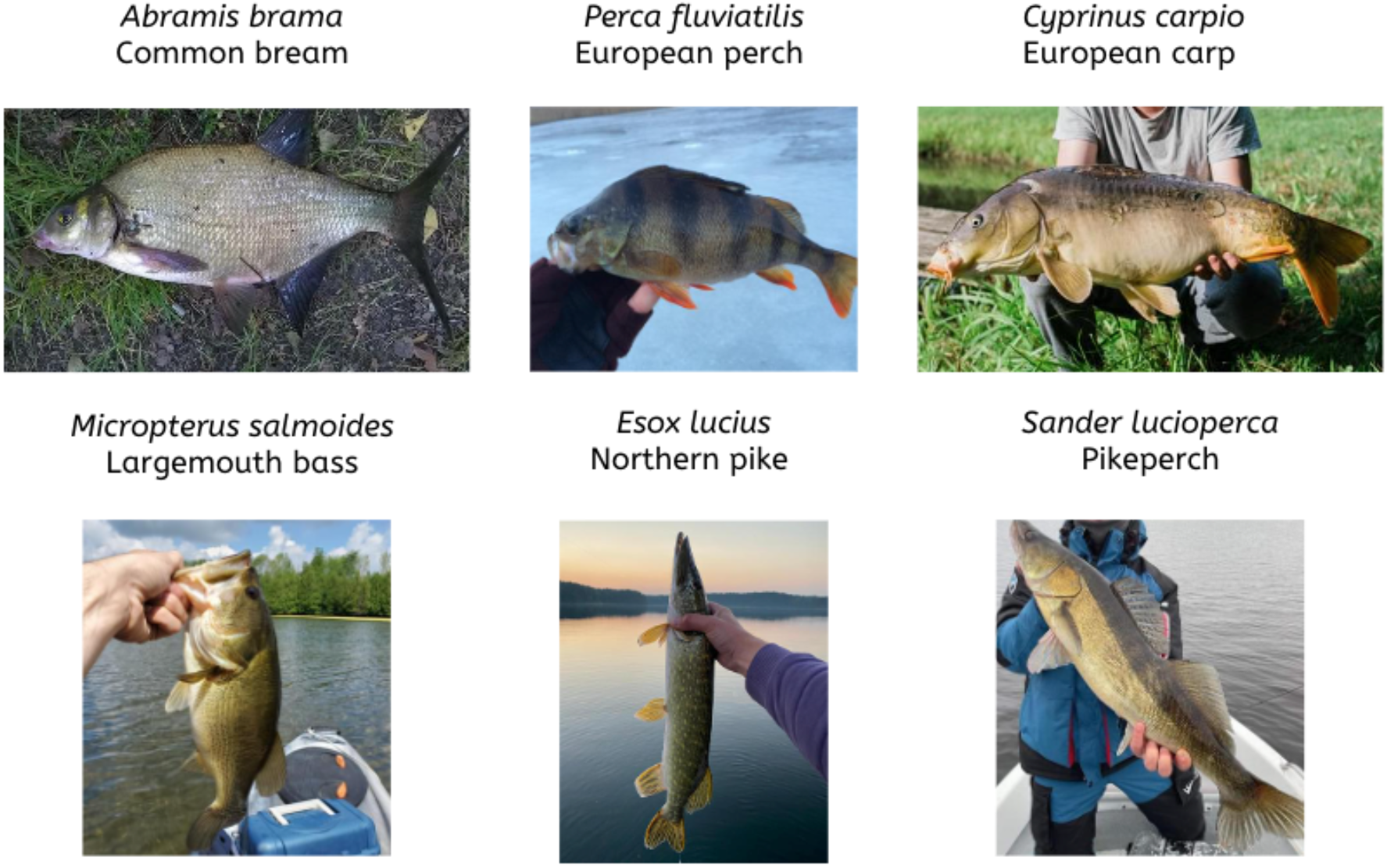
Example of images used for training the model, for each class (fish species)

The best performance for the image classification was achieved when using EfficientNet-Lite0 model architecture, a batch size of 32 and 20 epochs, with an overall accuracy of 0.91 and mean loss of 0.71 (Figure 3). Many classes had high precision values, although the precision for *Sander lucioperca* was quite low. From the confusion matrix (Figure 3), *Sander lucioperca* were commonly mistaken for *Esox lucius, Abramis brama* and *Perca fluviatilis*.

**Figure 3:**
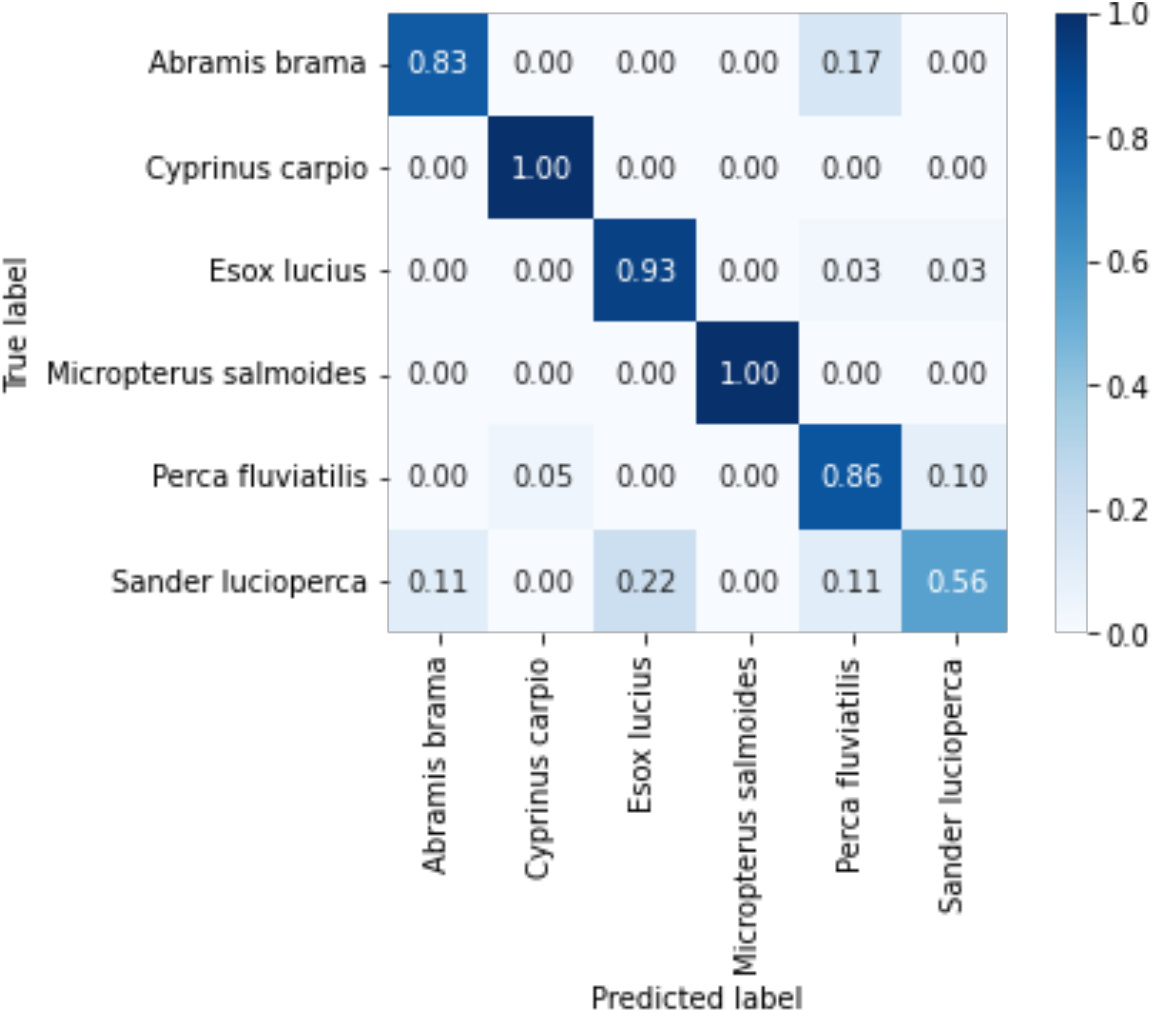
Confusion matrix of the image classification results with normalized, relative values of correct predictions for each species (i.e. precision) obtained when using EfficientNet-Lite0 model architecture, a batch size of 32 and 20 epochs.

For object detection, the best overall precision obtained was 0.48 when using EfficientDet-Lite0 model architecture, a batch size of 32 and 20 epochs (Table 4). Interestingly, only the class *Cyprinus carpio* exhibited high precision, indicating that the size of the training dataset by itself might not be the only indicator of model performance. Similarly to what other researchers found (e.g. Horn *et al*. 2017; Zheng *et al*. 2019), our results show that the classification method achieve higher overall performance when compared to the overall performance of the detection methods. Training the model with larger number of images, a balanced dataset and optimizing hyperparameters (such as epochs and batch size) may improve model performance.

**Table 3:** Sample sizes (images annotated) for both techniques: image classification and object detection.

| Species | Common name | Number of annotated images | Total number of images (after augmentation) |
| --- | --- | --- | --- |
| *Abramis brama* | Common bream | 111 | 333 |
| *Cyprinus carpio* | European carp | 377 | 1131 |
| *Esox lucius* | Northern pike | 420 | 1260 |
| *Micropterus salmoides* | Largemouth bass | 175 | 525 |
| *Perca fluviatilis* | European perch | 332 | 996 |
| *Sander lucioperca* | Pikeperch | 188 | 564 |

**Table 4:** Precision values for the best performing object detection model (model architecture = EfficientDet-Lite0, batch size = 32, epochs = 20)

| Species | Common name | Precision |
| --- | --- | --- |
| All (Average Precision) |  | 0.48 |
| *Abramis brama* | Common bream | 0.20 |
| *Cyprinus carpio* | European carp | 0.75 |
| *Esox lucius* | Northern pike | 0.55 |
| *Micropterus salmoides* | Largemouth bass | 0.53 |
| *Perca fluviatilis* | European perch | 0.31 |
| *Sander lucioperca* | Pikeperch | 0.53 |

## 4. Lessons learned, challenges and future applications

In our pilot case study we illustrate the scientific application, utility and potential for scalability of the framework presented here. The framework is flexible and can be customised and applied to a variety of image datasets and research questions. With a relatively small number of images per class (200-300) we demonstrate that a high performing model can be developed for a small number of classes by a small research group with few resources. Traditional deep learning-based approaches require training models on a dedicated server and high computational power to make the inference. To overcome this issue, this framework uses the Tensorflow Lite Model Maker library (Abadi et al. 2016a; b) and transfer learning which reduces the amount of training data required as well as model training time. In addition, this library is very flexible and new pre-trained models can be added by customising the library code.

While applying the framework to our pilot case study we have used a total of 59.39 GiB of cloud storage and computer resources included 16 vCPUs, 60 GB RAM and a Nvidia Tesla P4 GPU which resulted in a total of €117.20 (usage for three months). The cloud storage was used during the three month period and total expenses related to this service were €12.81. However, it is important to note that the costs were also kept low because we took advantage of free online tools with Python programming environments and free computing resources (Google colab) for most of exploratory work. Paid services of compute engine resources and notebooks were used only for intensive model runs.

We found that in the steps 2 and 3 the pre-trained model for detecting human faces or fish shapes were not always accurate. The pre-trained model *caffemodel* that was used to detect human faces sometimes failed to detect a face if it was not in a vertical position, as was the case in rotated images. False positives also occurred when the model placed bounding boxes on fish “faces”. This step could be improved by training a model to detect human faces using augmented data with transformations such as image rotation or vertical flip. Still, for images that need to be crowdsourced to public domains, for e.g., manual annotations or citizen science projects, the potential for sensitive data leakage must be carefully addressed. When it comes to detecting fish shapes and placing bounding boxes around them, in most cases the pretrained model *inception_resnet_v2* worked well, although often the box excluded small parts of the fish (usually end of the tail). However, when there were many overlapping fishes in the image, the model did not always detect all fishes in an image. In a few images, the pretrained model identified other object (e.g. shoes, boxes, etc.) as fish.

### Manual annotation is fast but might be even faster

Manual annotation of images was identified as an important challenge, where expert knowledge was crucial for correctly identifying fish species. The pre-annotation step (step 3) accelerated the process by automatically adding bounding boxes around fish, but images still needed to be individually assessed and identified. On average, for the six common and clearly distinct species, annotating one photo took about 2-3 seconds (although in some cases separating the common bream from other similar species took longer). In our case all annotated photos were divided into separate folders, depending on the month they were taken. Each folder contained c. 2000-2500 photos and up to half of them did not include any fish. In general, to review and annotate all the photos in the folder it took approximately 2-3 hours of intensive work by a highly skilled expert. This might be slowed down, depending on the speed of the computer and internet, or expertise level. As the model is developed and more species are added, new photos can be identified faster by applying the model to them first and then manually processing only those photos that had low classification score.

Citizen scientists can also supplement manual annotation of images for ML projects. For example, Gundelund *et al*. (2021) show that citizen scientists can estimate fisheries metrics and identify species with results comparable to surveys by scientists. If images can be shared publicly, crowdsourcing citizen science platforms Zooniverse (https://www.zooniverse.org/) could speed up the annotation and its accuracy. For example, Anton et al. (2021) used the platform to engage many citizen scientists to efficiently and accurately annotate data from underwater footage to detect cold water corals. To ensure higher accuracy the authors used repeated annotations, i.e. each video clip was annotated by eight citizen scientists and an agreement threshold of 80% was used.

Like other researchers (e.g. Lekunberri *et al*. 2022), we found that image augmentation through rotation and flips improved the image classification model performance (overall accuracy increased by 10%). The augmentation was easy, fast and straightforward and we recommend using it in most image classification and object detection applications.

In our pilot case study, the process of model training using the classification technique took 6903 seconds (approximately 2 hours), while using the object detection technique it took 3906 seconds (approximately 1 hour) for the same number of images (n=4809) batches (n=32) and epochs (n=20) but a different model architecture (EfficientNet-Lite0 and EfficientDet-Lite0 respectively). The time required for the training phase of a model is an important consideration when choosing which computer vision technique to use. For the same amount of data and hyperparameter space (batch size and epochs) image classification can require more time to train a model, when comparing to the object detection technique. However, to achieve higher performance, object detection methods might need larger amount of data and more time for hyperparameters optimization. In addition, the time consuming process of manual annotation of images for object detection is also important to consider.

In fisheries contexts, the majority of image processing and classification models aim to automatically identify fish species and are developed at the regional level, and are often fisheries-specific (Lekunberri *et al*. 2022; Ovalle *et al*. 2022; Palmer *et al*. 2022). Even though groups use different techniques (such as image classification, object detection and segmentation) these individual (and regional) models could be combined into a global, hierarchical classification framework for automatically identifying fish species worldwide. The process of combining different machine learning models is called ensemble learning. Usually, an ensemble classification model consists of two steps: (1) generating classification results using multiple weak classifiers, and (2) integrating multiple results into a consistency function to get the final result with voting schemes (Dong *et al*. 2020). There are different methods of ensemble learning with their own advantages and disadvantages (reviewed in Dong *et al*. 2020) and this area of research is rapidly evolving. However, ensemble learning has already been recognised to improve the performance of individual models and building a new model by ensemble learning requires less time, data and computational resources than training a new model with all the data combined.

Global open-access machine learning models would have a range of applications for research and fisheries management. Platforms such as iNaturalist and Fishbase.org could benefit from fisheries-specific models or models developed at regional levels which can be combined by ensemble learning. Data collection can be further accelerated as tasks such as automatic fish identification, size estimation or sex determination can be speeded up with model predictions therefore helping citizen scientists with the process of metadata entry. In return, research projects and management efforts could take advantage of these platforms and data.

## Data availability

All steps described and scripts used are supported by a freely available code library, deposited in github.com/FishSizeProject

## References

Abadi M, Agarwal A, Barham P, Brevdo E, Chen Z, Citro C, Corrado GS, Davis A, Dean J, Devin M, Ghemawat S, Goodfellow I, Harp A, Irving G, Isard M, Jia Y, Jozefowicz R, Kaiser L, Kudlur M et al. (2016a) TensorFlow: Large-Scale Machine Learning on Heterogeneous Distributed Systems. arXiv.

Abadi M, Barham P, Chen J, Chen Z, Davis A, et al. (2016b) TensorFlow.js. In: Proceedings of the 12th USENIX Symposium on Operating Systems Design and Implementation

Anton V, Germishuys J, Bergström P, Lindegarth M, Obst M (2021) An open-source, citizen science and machine learning approach to analyse subsea movies. Biodiversity Data Journal, 9, 1–14.

Bengio Y, Bastien F, Bergeron A, Boulanger-Lewandowski N, Breuel T, Chherawala Y, Cisse M, Côté M, Erhan D, Eustache J, Glorot X, Muller X, Lebeuf SP, Pascanu R, Rifai S, Savard F, Sicard G (2011) Deep learners benefit more from out-of-distribution examples. Proceedings of the Fourteenth International Conference on Artificial Intelligence and Statistics, 15, 164–172.

Bisong E (2019) Building Machine Learning and Deep Learning Models on Google Cloud Platform. In: Building Machine Learning and Deep Learning Models on Google Cloud Platform, pp. 59–64. Berkeley, CA.

Buslaev A, Iglovikov VI, Khvedchenya E, Parinov A, Druzhinin M, Kalinin AA (2020) Albumentations: Fast and flexible image augmentations. Information, 11, 1–20.

Costello C, Ovando D, Hilborn R, Gaines SD, Deschenes O, Lester SE (2020) Status and Solutions for the World’s Unassessed Fisheries. Science, 338, 517–521.

Dickinson JL, Zuckerberg B, Bonter DN (2010) Citizen science as an ecological research tool: Challenges and benefits. Annual Review of Ecology, Evolution, and Systematics, 41, 149–172.

Dong X, Yu Z, Cao W, Shi Y, Ma Q (2020) A survey on ensemble learning. Frontiers of Computer Science, 14, 241–258.

Dutta A, Zisserman A (2019) The VIA annotation software for images, audio and video. MM 2019 - Proceedings of the 27th ACM International Conference on Multimedia, 2276–2279.

Ebrahimi SH, Ossewaarde M, Need A (2021) Smart fishery: A systematic review and research agenda for sustainable fisheries in the age of ai. Sustainability (Switzerland), 13.

Gundelund C, Venturelli P, Hartill BW, Hyder K, Olesen HJ, Skov C (2021) Evaluation of a citizen science platform for collecting fisheries data from coastal sea trout anglers. Canadian Journal of Fisheries and Aquatic Sciences, 78, 1576–1585.

Harris D, Johnston D, Yeoh D (2021) More for less: Citizen science supporting the management of small-scale recreational fisheries. Regional Studies in Marine Science, 48, 102047.

He K, Zhang X, Ren S, Sun J (2016) Deep residual learning for image recognition. Proceedings of the IEEE Computer Society Conference on Computer Vision and Pattern Recognition, 770–778.

Hendrycks D, Mu N, Cubuk ED, Zoph B, Gilmer J, Lakshminarayanan B (2020) AugMix: A Simple Data Processing Method to Improve Robustness and Uncertainty. In: Proceedings of the International Conference on Learning Representations (ICLR), pp. 1–15.

Hernández-García A, König P (2018) Further advantages of data augmentation on convolutional neural networks. In: Lecture Notes in Computer Science (including subseries Lecture Notes in Artificial Intelligence and Lecture Notes in Bioinformatics), pp. 95–103.

Horn G Van, Mac O, Shepard A, Adam H, Song Y, Cui Y, Sun C, Perona P, Belongie S (2017) The iNaturalist Species Classification and Detection Dataset - Supplementary Material. Computer Vision Foundation, 4–6.

Kluyver T, Ragan-Kelley B, Pérez F, Granger B, Bussonnier M, Frederic J, Kelley K, Hamrick J, Grout J, Corlay S, Ivanov P, Avila D, Abdalla S, Willing C (2016) Jupyter Notebooks—a publishing format for reproducible computational workflows. Positioning and Power in Academic Publishing: Players, Agents and Agendas - Proceedings of the 20th International Conference on Electronic Publishing, ELPUB 2016, 87–90.

Kuznetsova A, Rom H, Alldrin N, Uijlings J, Krasin I, Pont-Tuset J, Kamali S, Popov S, Malloci M, Kolesnikov A, Duerig T, Ferrari V (2020) The Open Images Dataset V4: Unified Image Classification, Object Detection, and Visual Relationship Detection at Scale. International Journal of Computer Vision, 128, 1956–1981.

Lekunberri X, Ruiz J, Quincoces I, Dornaika F, Arganda-Carreras I, Fernandes JA (2022) Identification and measurement of tropical tuna species in purse seiner catches using computer vision and deep learning. Ecological Informatics, 67.

Liu S, Papailiopoulos D, Achlioptas D (2020) Bad global minima exist and SGD can reach them. Advances in Neural Information Processing Systems, 2020-Decem.

Meirelles K, Freire F, Belhabib D, Espedido JC, Hood L, Kleisner KM, Lam VWL, Machado ML, Mendonça JT, Meeuwig JJ, Moro PS, Motta FS, Palomares MD, Smith N, Teh L, Zeller D, Zylich K, Pauly D, Purcell SW et al. (2020) Estimating Global Catches of Marine Recreational Fisheries. Frontiers in Marine Science, 7, 1–18.

Mikołajczyk A, Grochowski M (2018) Data augmentation for improving deep learning in image classification problem. In: 2018 International Interdisciplinary PhD Workshop, IIPhDW 2018, pp. 117–122.

Mohri M, Rostamizadeh A, Talwalkar A (2012) Foundations of Machine Learning. MIT Press, Cambridge, MA.

Ovalle JC, Vilas C, Antelo LT (2022) On the use of deep learning for fish species recognition and quantification on board fishing vessels. Marine Policy, 139, 105015.

Palmer M, Álvarez-Ellacuría A, Moltó V, Catalán IA (2022) Automatic, operational, high-resolution monitoring of fish length and catch numbers from landings using deep learning. Fisheries Research, 246.

Ponnusamy A (2018) cvlib - high level Computer Vision library for Python.

Prince SJD (2012) Computer vision: Models, learning and inference. Cambridge University Press.

Sandler M, Howard A, Zhu M, Zhmoginov A, Chen LC (2018) MobileNetV2: Inverted Residuals and Linear Bottlenecks. Proceedings of the IEEE Computer Society Conference on Computer Vision and Pattern Recognition, 4510–4520.

dos Santos AA, Gonçalves WN (2019) Improving Pantanal fish species recognition through taxonomic ranks in convolutional neural networks. Ecological Informatics, 53, 100977.

Shorten C, Khoshgoftaar TM (2019) A survey on Image Data Augmentation for Deep Learning. Journal of Big Data, 6.

Tan M, Le Q V. (2019) EfficientNet: Rethinking model scaling for convolutional neural networks. 36th International Conference on Machine Learning, ICML 2019, 10691–10700.

Tan M, Pang R, Le Q V. (2020) EfficientDet: Scalable and efficient object detection. Proceedings of the IEEE Computer Society Conference on Computer Vision and Pattern Recognition, 10778–10787.

Venturelli PA, Hyder K, Skov C (2017) Angler apps as a source of recreational fisheries data: opportunities, challenges and proposed standards. Fish and Fisheries, 18, 578–595.

Zheng YY, Kong JL, Jin XB, Wang XY, Su TL, Zuo M (2019) Cropdeep: The crop vision dataset for deep-learning-based classification and detection in precision agriculture. Sensors (Switzerland), 19.

